# Cross-reactivity of SARS-CoV-2-specific T cells against tumor-associated antigens via molecular mimicry

**DOI:** 10.64898/2026.02.16.706083

**Authors:** Concetta Ragone, Angela Mauriello, Beatrice Cavalluzzo, Simona Mangano, Biancamaria Cembrola, Noemi Ciotola, Maria Tagliamonte, Luigi Buonaguro

**Affiliations:** Innovative Immunological Models Unit, Istituto Nazionale Tumori - IRCCS - “Fond G. Pascale”, Naples - ITALY

## Abstract

**BACKGROUND:** We have recently described SARS-COV-2 antigens showing sequence and conformational homology to tumor associated antigens (TAAs). Moreover, cross-reactive T cells have been identified in individuals either infected by the SARS-CoV-2 virus or vaccinated with the BNT162b2 preventive vaccine. In the present study, we analyzed the specific cross-binding TCRs by single cell RNA TCR sequencing.

**METHODS AND RESULTS:** The paired SARS-CoV-2 epitope LLLDDFVEI (VIR) and the PRDX5 tumor associated antigen LLLDDLLVS (TAA) were selected to elicit cross-reacting T cells *ex vivo*. PBMCs from 5 healthy individuals were cultured for 10 days with 10 ug every 3 days of one of the two peptides and cells were selected for single cell RNA TCR sequencing. Results in CD8^+^ T Effector cells (T_TE_) showed the amplification or the *de novo* identification of a handful number of TRAV/TRBV genes and of CDR3αβ motifs upon treatment *ex vivo* with both epitopes, which are specific for each subject in the analysis. The very same clonotypes were identified also in the CD8^+^ T proliferating subset, confirming that both epitopes induced a highly activated and plastic state. Conformational prediction analyses of pMHC-TCR complexes showed perfect structural overlap, supporting the functional cross-reaction of CD8+ T cells with both the viral and the tumor antigens.

**CONCLUSIONS:** Our results describe for the first time the TCR CDR3αβ motifs amplified or *de novo* expanded by induction with a viral antigen showing a molecular mimicry with a tumor antigen. They are strictly individual and do not match with any motif in the publicly available TCR repository. However, considering the significant degeneracy in the TCR binding to the same epitope, the finding of identical TCR CDR3αβ motifs elicited by two homologous epitopes is of the highest functional relevance. Such results provide a clear experimental validation proof that microbial epitopes mimicking TAAs can be used to develop off-the-shelf preventive/therapeutic vaccine formulations. Indeed, such non-self antigens are much stronger immunogens and may elicit a potent cross-reacting anti-cancer T cell response.

## INTRODUCTION

The success of therapeutic cancer vaccines, mostly based on wild-type tumor associated antigens (TAAs), has been limited largely because they are self-antigens and suffer from immune tolerance^1^. On the contrary, cancer vaccines based on personalized tumor specific neoantigens (TSAs) hold great promise but, at the present stage, significant obstacles pertaining their design and the tumour environment remain to be overcome^2^. In such a framework, aiming at developing off-the-shelf therapeutic cancer vaccines based on shared over-expressed TAAs, we need to overcome their intrinsic low immunogenicity. The exploitation of the molecular mimicry for developing effective cancer vaccines with microbe-derived non-self immunogenic antigens (MAAs) might represent the turning point. They would elicit a potent T cell response which will cross-react against highly similar TAAs^3,4^. Indeed, the T cell immunity incorporates an extremely high level of receptor degeneracy, enabling each TCR to recognize multiple peptides^5^.

In the recent years, several MAAs have been reported sharing sequence or structural similarities with either well characterized TAAs^6-11^. Likewise, similarities have been shown reported between MAAs and “non canonical” TAAs^12,13^. Cross-reactive T cells have been identified in healthy individuals or in subjects undergoing natural infection and/or vaccination (i.e. HIV-1, SARS-CoV-2)^10,14^.

HLA-A^*^02:01+ healthy donors and melanoma patients have been shown to have circulating CD8^+^ T cells reactive to the Melan-A/MART-1_27-35_ showing a molecular mimicry with ubiquitous herpes simplex virus 1 (HSV1) and HSV2 derived epitopes^15,16^.

Tumor-infiltrating lymphocytes (TILs) have been identified in human lung cancer tissues cross-reacting with an antigen derived from the Epstein-Barr virus (EBV) latent membrane protein 2a (LMP2a) and a highly homologous antigen derived from the TMEM161A protein^17^. Moreover, we have recently shown that peptides derived from flagellin-related mimic the structure of melanoma TAAs and a reactivity against the bacterial peptides is preferentially detected in PBMCs and in TILs from patients responding to Immune CheckPoint inhibitors (ICI_S_)^18^.

Moreover, anectodical observations have reported a significant tumor burden reduction in 3 patients affected by metastatic colorectal cancer (mCRC), during infection by severe acute respiratory syndrome coronavirus 2 (SARS-Cov-2)^19^.

All such experimental evidences have not been supported yet by the description of TCR CDR3αβ motifs selectively elicited by the homologous MAA/TAA pairs.

The aim of the present study was to fill this gap assessing the TCR sequences specifically elicited *ex vivo* by the peptide pair SARS-CoV-2 LLLDDFVEI and the PRDX5 TAA LLLDDLLVS^10^. The outcome of the scRNA TCR sequencing analysis has identified a number of TCR CDR3αβ motifs amplified or *de novo* expanded following the treatment with either peptides when compared to the baseline. Even more interestingly, a set of such motifs are found in samples following the treatment with both peptides.

This represents the first documentation of TCR CDR3αβ motifs involved in the recognition of a viral antigens showing the molecular mimicry with a TAA, providing an additional key piece of information in the exploitation of such a strategy for developing cancer vaccines targeting shared TAAs.

## MATERIALS AND METHODS

### PBMCs isolation and ex vivo stimulation

Peripheral blood was obtained by venipuncture from five HLA-A^*^02:01–positive healty donors enrolled at the National Cancer Institute “Pascale” in Naples, ITALY. Fresh human PBMCs were isolated by Ficoll-Hypaque density gradient centrifugation and cultured in ImmunoCult™-XF T Cell Expansion Medium (STEMCELL Technologies) supplemented with 2 mM l-glutamine (Sigma), 10% fetal bovine serum (Life Technologies) and 2%penicillin/streptomycin (5000 I.U./5 mg/ml, MP Biomedicals). Viral peptide (LLLDDFVEI) or TAA peptide ((LLLDDLLVS) were added at a final concentration of 10 ug/mL every 3 days. Following 10 days of ex vivo stimulation, CD8^+^ T cells were isolated through human CD8^+^ T Cell Isolation Kit (Miltenyi Biotec) and subjected to single cell RNA TCR sequencing.

### Library Preparation

Single cells were suspended in phosphate‐buffered saline containing 0.04% bovine serum albumin, filtered using a 40 μm cell strainer (Biologix), and their concentration was evaluated using the LUNA-II™ Automated Cell Counter (Logos Biosystems). The cell suspension was loaded onto the Chromium Single Cell G Chip Kit (10x Genomics) and processed on the Chromium X Single Cell Controller to generate single-cell gel bead-in-emulsions (GEMs), according to the manufacturer’s instructions.

Single-cell gene expression and T-cell receptor (TCR) libraries were generated using the Chromium Single Cell 5′ Library & Gel Bead Kit in combination with the Chromium Single Cell V(D)J Enrichment Kit, enabling the simultaneous capture of whole-transcriptome profiles and full-length TCR α and β chain sequences from the same cells. Following reverse transcription and cDNA amplification, gene expression and TCR libraries were constructed according to the manufacturer’s protocol. cDNA quality was assessed using High Sensitivity D5000 ScreenTape on an Agilent 4200 TapeStation system (Agilent Technologies), while final library quality was evaluated using High Sensitivity DNA D1000 ScreenTape (Agilent Technologies). Finally, libraries were sequenced on a NovaSeq X platform (Illumina) according to the manufacturer’s specifications.

### Bioinformatics workflow

Illumina NovaSeq 6000 base call (BCL) files were converted into fastq files through bclconvert. Mapping on reference genome, counting per genes and TCR detection were performed using Cell Ranger multi1 (v. 8.0.1), default setting. Annotation profile was performed using Celltypist tool (https://www.celltypist.org/), Healthy_COVID19_PBMC model. Downstream analyses were performed using Seurat and reticulate package in R (version ≥4.2), obtaining a data frame with UMAP coordinates, annotations, and cell barcodes. For each sample, unique cell barcodes corresponding to CD8 T cells were used to compute cell-type composition. Counts and relative frequencies (percentages) were calculated for each CD8 T-cell subtype (CD8.TE, CD8.EM, CD8.Naive, CD8.Prolif). Bar plots were generated using ggplot2 package in R in order to visualize the relative abundance of CD8 T-cell subtypes across samples and treatments. Samples were grouped by patient prefix and treatment condition (Untreated, ORF, TAA). Percentages were displayed using stacked bar plots, with consistent color coding across all figures. UMAP visualizations were generated for CD8 T cells only, using ggplot2 package in R. To improve rendering performance for large datasets, point layers were rasterized. Cluster labels were positioned using a centroid-based approach: for each CD8 subtype, the cell with the minimal summed Euclidean distance to all other cells in the same cluster was selected as the label anchor.

### T cell receptor analysis

Data from scRNA sequencing in each donor were analyzed from cells untreated (UNT) as well as treated *ex vivo* with either the viral antigen (VIR) or the tumor antigen (TAA). For the endogenous repertoires, only expression levels from the UNT samples were taken into consideration. The T cell clones differentially expressed in the VIR and TAA samples were identified by comparison with values in the UNT samples. Clones were selected based on the parameter of an increased expression in both VIR and TAA samples vs. UNT samples (>1.5 fold).

### Analysis of CDR3αβ motifs

The CDR3αβ clonotypes expanded after induction with both VIR and TAA in the present study were searched in the VDJdb database https://www.vdjdb.com/. CDR3αβ motifs were selected and piled up to generate sequence logos as are a graphical representation of the information content stored in a multiple sequence alignment (MSA) and provide a compact and highly intuitive representation of the position-specific amino acid composition of binding motifs, active sites, etc. in biological sequences. Accurate generation of sequence logos is often compromised by sequence redundancy and low number of observations. Moreover, most methods available for sequence logo generation focus on displaying the position-specific enrichment of amino acids, discarding the equally valuable information related to amino acid depletion. Seq2logo aims at resolving these issues allowing the user to include sequence weighting to correct for data redundancy, pseudo counts to correct for low number of observations and different logotype representations each capturing different aspects related to amino acid enrichment and depletion. Besides allowing input in the format of peptides and MSA, Seq2Logo accepts input as Blast sequence profiles, providing easy access for non-expert end-users to characterize and identify functionally conserved/variable amino acids in any given protein of interest. The output from the server is a sequence logo and a PSSM. Seq2Logo is available at http://www.cbs.dtu.dk/biotools/Seq2Logo.

### Structural modeling tool of TCR-pMHC class I complexes

Three-dimensional model of the TCR-pMHC complexes, has been generated with the *TCRpMHCmodels 1.0* tool (https://services.healthtech.dtu.dk/services/TCRpMHCmodels-1.0/) including the peptide sequences of TCR alpha chain, TCR beta chain, peptide and MHC. Upon submission, this tool automatically selects the best templates and generates models of the TCR-pMHC complex, with a median Cα RMSD of 2.3Å. The 3D structures were generated using the Molsoft ICM (http://www.molsoft.com/, version 3.8-7d) software.

## RESULTS

### Expansion of proliferating and effector memory CD8^+^ T cells

To profile the phenotypic and functional states of T cell clones, we constructed an atlas comprising ∼100,000 CD8^+^ T cells collected from 5 donors. For each donor, we analyzed 3 samples: untreated (UNT), treated with the viral peptide (VIR) or with the tumor antigen (TAA).

Cell annotation showed a large degree of variation in CD8^+^ T cell subsets composition among donors individuals, with an overall expansion of effector, effector memory and proliferating T cells clusters. Indeed, UMAP projection of CD8^+^ T cells from all the 5 donors revealed a consistent pattern of CD8^+^ T-cell distinct differentiation states across conditions (UNT vs. VIR vs. TAA). The untreated sample (UNT) showed well-segregated naïve, effector memory, terminal effector and proliferative populations, consistent with a basal differentiation state (Fig. 1). On the contrary, the sample treated with the viral peptide (VIR) samples displayed marked expansion of proliferating and effector memory CD8^+^ T cells with increased overlap between subsets, consistent with a highly activated and plastic state. In parallel, the sample treated with the tumor antigen peptide (TAA) exhibited reduced proliferative activity and a predominance of effector and terminal effector phenotypes, suggesting a more differentiated CD8^+^ T-cell response. These patterns are consistent with condition-specific modulation of CD8^+^ T-cell differentiation and activation states.

**Figure 1.**
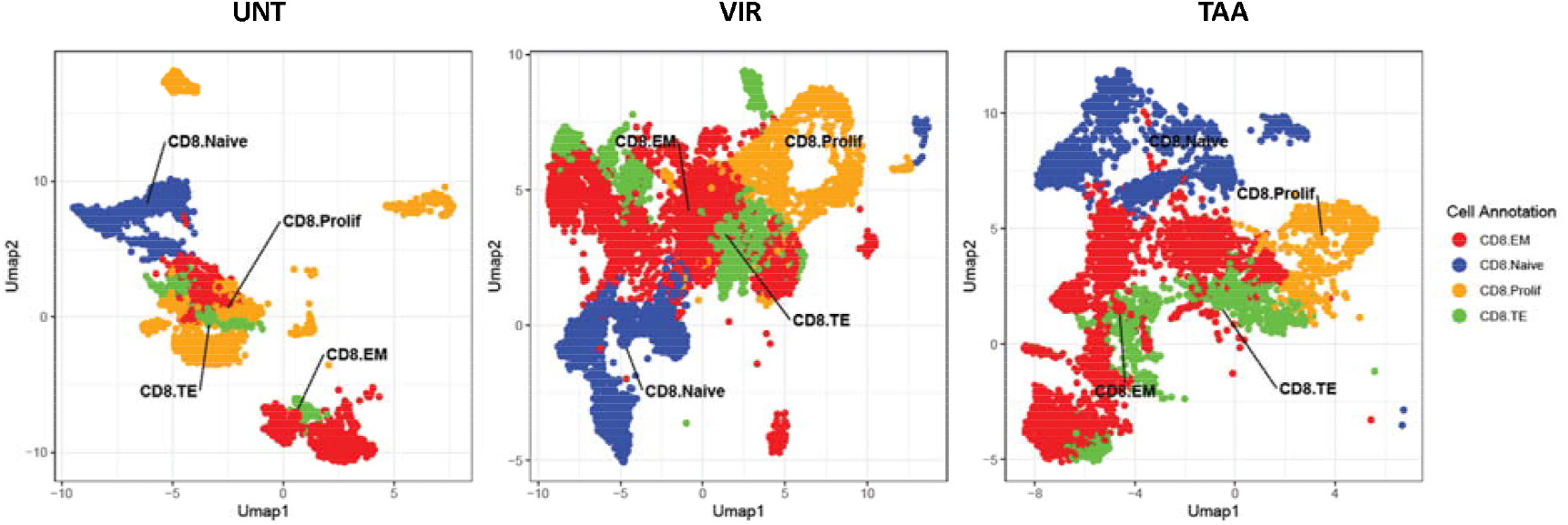
Uniform Manifold Approximation and Projection (UMAP) of single-cell transcriptomic profiles of CD8^+^ T cells from a single donor in UNT, VIR and TAA conditions. Each point represents an individual CD8^+^ T cell, colored according to cell-state annotation: naïve (CD8.Naive, blue), effector memory (CD8.EM, red), terminal effector (CD8.TE, green), and proliferating (CD8.Prolif, orange).

### TCRαβ endogenous repertoires

In order to assess the endogenous basal T cell receptor (TCR) repertoire in each subject analyzed in the present study, a quantification of the TCRαβ chains of was assessed in the CD8^+^ T Effector cells (T_TE_). Indeed, they are the subset actively responding to a stimulus in order to eliminate infected cells and protect the host from severe infection^20^.

The results from single cell TCR RNA sequencing analysis showed individual unique patterns of the T-cell receptor αβ variable (TRAV & TRBV) clones, with few of them prevalently expanded in each subject. The latter do not show a clear overlapping between subjects, with a single exception in the TRAV (e.g. TRAV8-3) and few exceptions in the TRBV (e.g. TRBV5-1; TRBV19; TRBV7-9) (Fig. 2A).

**Fig. 2A.**
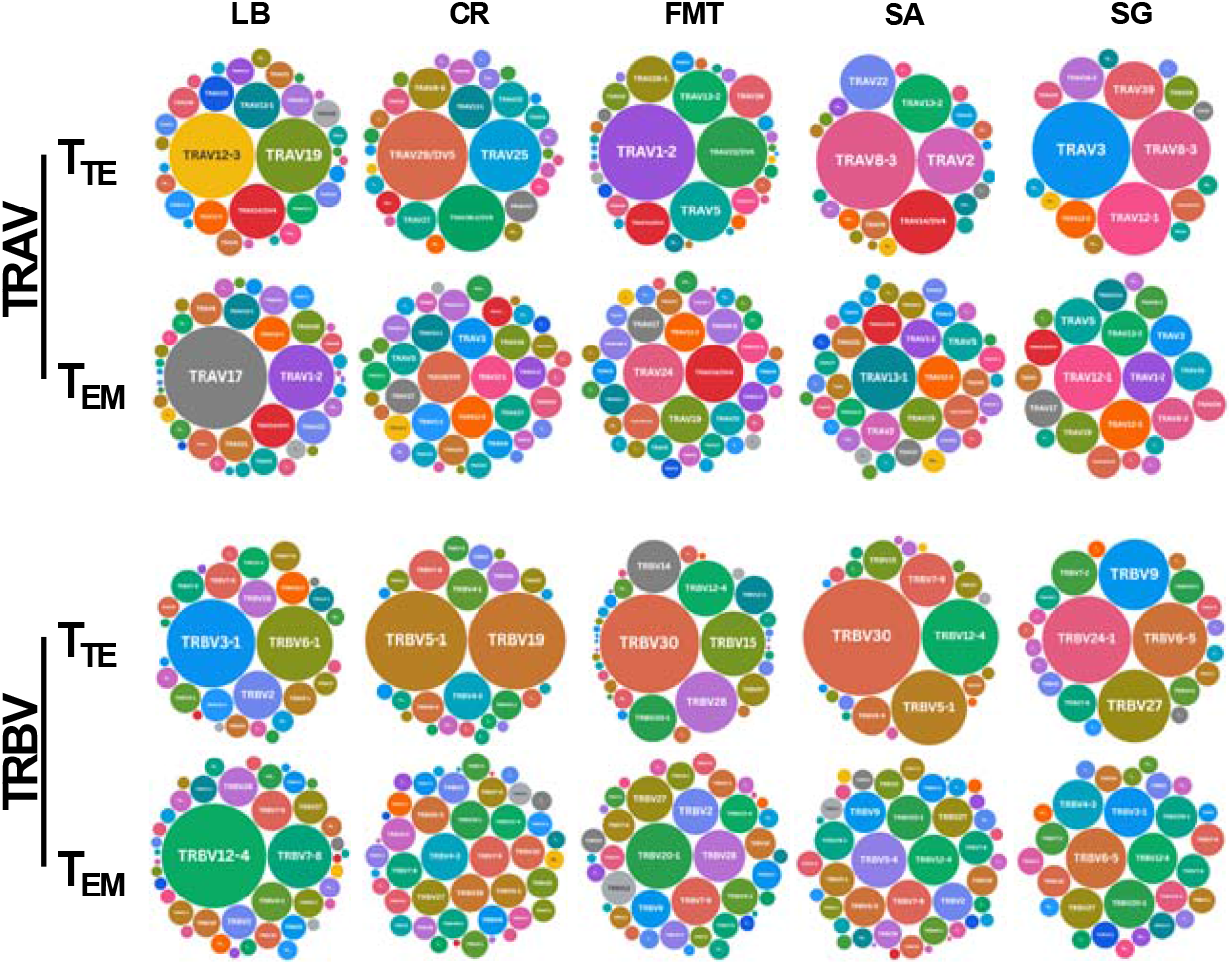
TRAV and TRBV endogenous repertoire. The indipendent TRAV and TRBV repertoire in each donor is represented in CD8^+^ T Effector cells (T_TE_) and Effector Memory (T_EM_) subsets. Clones’ size are indicated by symbol size.

The subsequent step was to assess whether the clones expanded in the CD8^+^ T Effector cells (T_TE_) were also predominant in the pool of Effector Memory T cells (T_EM_). The results showed a significantly different pattern, characterized by a general lack of highly prevalent clones over the background. Only the subject LB showed prevalent TRAV (e.g. TRAV17 and TRAV1-2) and TRBV (TRBV 12-4 and TRBV 7-8) clones in Effector Memory T cells (T_EM_). However, they were different from the ones expanded in the T Effector cells (T_TE_). The same lack of correlation between the clones expanded in the two CD8^+^ T cell subtypes was observed in all other subjects (Fig. 2A).

Furthermore, the same data were analyzed by generating unsupervised heatmaps. The results showed two clear clusters formed by the data from the CD8^+^ T_TE_ or the CD8^+^ T_EM_ subsets. Interestingly, some of the prevalent TRAV and TRBV molecules in the T_TE_ subset derive from the low prevalent ones in the T_EM_ subset (Fig. 2B). All the subsequent analyses were performed on the CD8^+^ T_TE_ subset.

**Fig. 2B.**
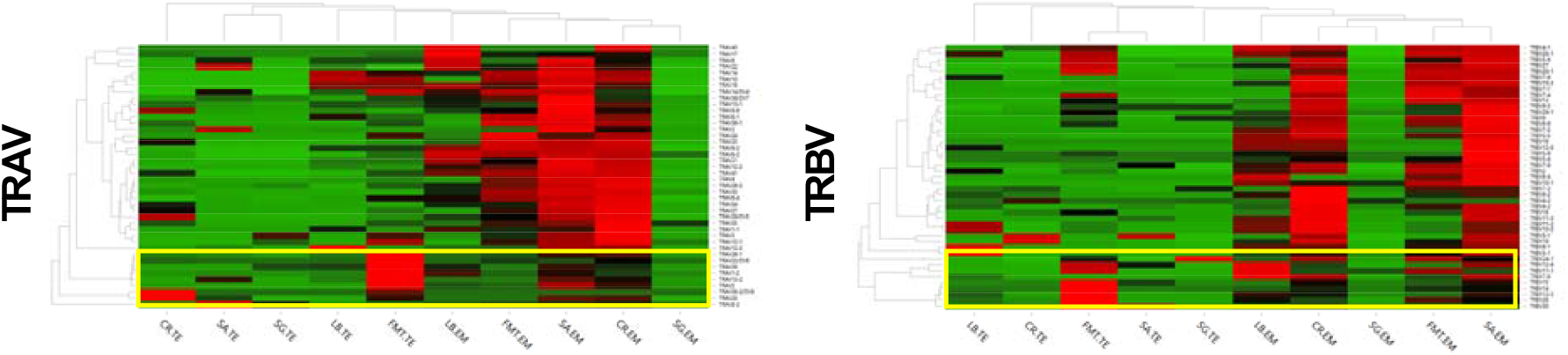
TRAV and TRBV endogenous repertoire. The endogenous expression levels of both TRAV and TRBV molecules are clustered according to the T cell subsets. The yellow box highlights the molecules significantly more prevalent in the T Effector cells (T_TE_) than in the Effector Memory (T_EM_) subset.

### TRAV & TRBV patterns elicited by VIR and TAA antigens in the CD8^+^ T_TE_ subset

In order to assess the TRAV & TRBV patterns expanded by the VIR or TAA antigen, scRNA TCR sequencing analyses were performed on PBMCs after *ex vivo* treatment with the peptides. The results showed that the expression levels of TRAV and TRBV clones in samples derived after culturing with VIR or TAA peptide, form independent donor-specific clusters (Fig. 3A-B).

**Fig. 3.**
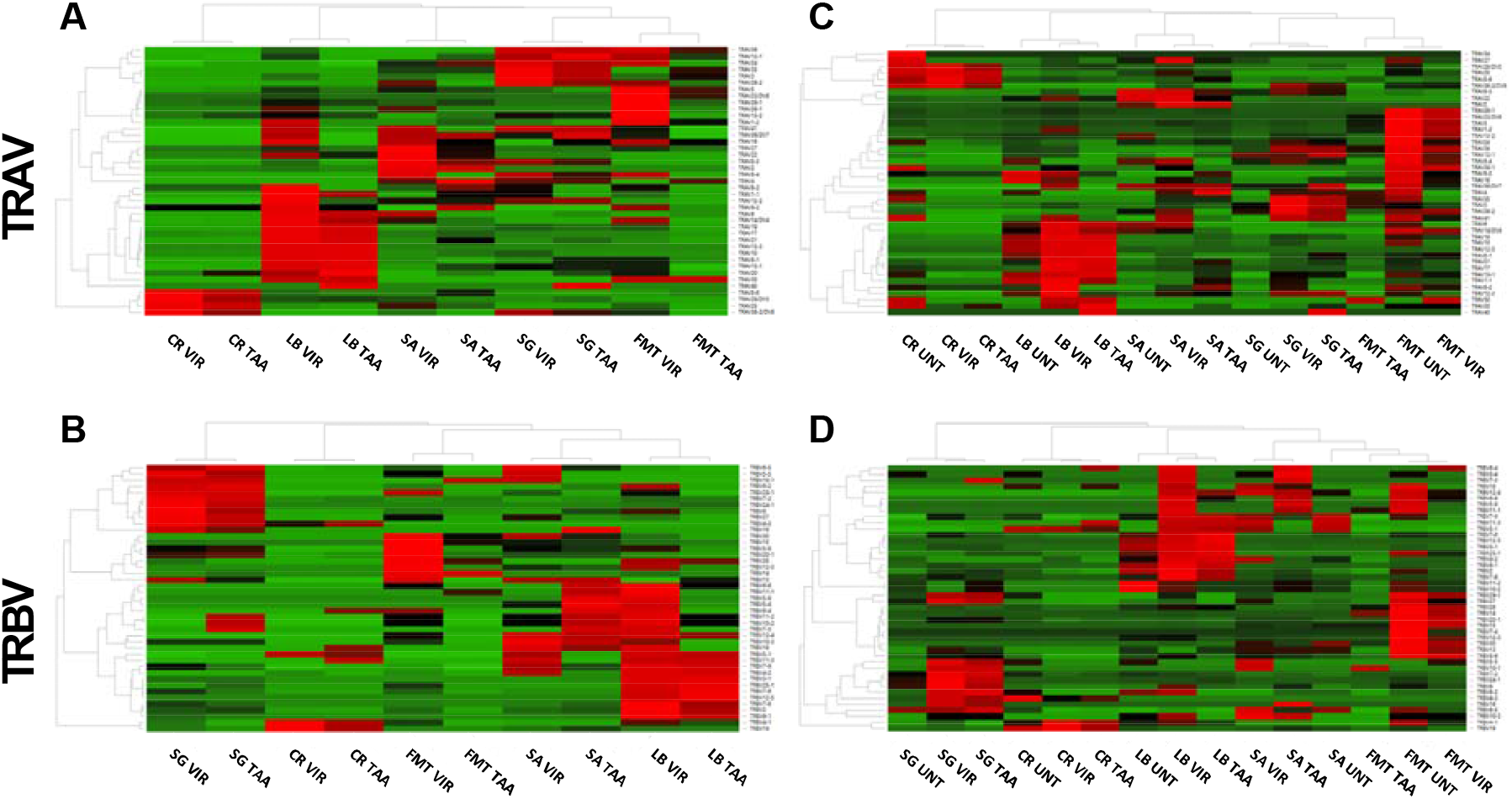
TRAV and TRBV molecules expanded ex vivo by epitopes CD8^+^ T_TE_ subset. The expression levels of both TRAV and TRBV molecules expanded ex vivo by both VIR and TAA epitope are clustered according to the donor (A). The same clustering is confirmed also when the sample untreated (UNT) are included in the heatmap analysis.

The same analysis performed including also the data from the untreated samples (UNT) showed that the three samples from each donor (UNT, VIR, TAA) form independent donor-specific clusters. It was confirmed that the samples after treatment with VIR or TAA show more similarity compared to the untreated sample (Fig. 3C-D). The only exception is the FMT donor. Such results suggest that both peptides induce similar expression levels in each donor, which are distinctive from each other.

A significant number of TRAV (nr. 14) and TRBV (nr. 10) clones show reactivity with either the VIR or the TAA antigen in at least 4 of the 5 subjects of the present analysis. However, considering as relevant a minimum of 5 clones increased after expansion with both antigens, 21 TRAV and TRBV are identified at least in a single donor and 7 at least in 2 donors (Tables 1 and 2).

**TABLE 1.** Number of TRAV clonotypes identified in each donor.

|  | LB |  |  | CR |  |  | FMT |  |  | SA |  |  | SG |  |  | A | B | C | D |
| --- | --- | --- | --- | --- | --- | --- | --- | --- | --- | --- | --- | --- | --- | --- | --- | --- | --- | --- | --- |
| TRAV | UNT | VIR | TAA | UNT | VIR | TAA | UNT | VIR | TAA | UNT | VIR | TAA | UNT | VIR | TAA |  |  |  |  |
| 10 | 13 | 26 | 26 | 0 | 0 | 0 | 10 | 0 | 0 | 0 | 0 | 1 | 0 | 2 | 0 |  |  | XXX |  |
| 1-1 | 9 | 16 | 6 | 0 | 0 | 0 | 1 | 1 | 0 | 0 | 0 | 3 | 0 | 3 | 0 |  |  |  |  |
| 1-2 | 8 | 116 | 5 | 0 | 0 | 0 | 455 | 181 | 35 | 4 | 5 | 16 | 1 | 9 | 3 | X | XX | XXX |  |
| 12-1 | 5 | 5 | 1 | 3 | 1 | 0 | 39 | 13 | 5 | 3 | 1 | 4 | 14 | 13 | 24 | X |  |  |  |
| 12-2 | 13 | 37 | 6 | 3 | 0 | 0 | 11 | 4 | 2 | 5 | 11 | 9 | 4 | 15 | 16 | X | XX | XXX | XXXX |
| 12-3 | 76 | 149 | 160 | 1 | 0 | 0 | 3 | 0 | 0 | 4 | 1 | 2 | 1 | 0 | 0 |  |  | XXX |  |
| 13-1 | 22 | 36 | 40 | 16 | 3 | 5 | 18 | 13 | 1 | 6 | 7 | 11 | 1 | 14 | 13 | X | XX | XXX | XXXX |
| 13-2 | 10 | 9 | 6 | 0 | 4 | 6 | 143 | 43 | 1 | 37 | 1 | 4 | 0 | 10 | 8 | X |  | XXX |  |
| 14/DV4 | 32 | 116 | 56 | 5 | 0 | 2 | 96 | 57 | 1 | 46 | 35 | 12 | 0 | 2 | 2 | X |  | XXX |  |
| 16 | 8 | 3 | 0 | 0 | 0 | 0 | 8 | 2 | 1 | 0 | 2 | 1 | 0 | 0 | 1 |  |  |  |  |
| 17 | 0 | 139 | 101 | 9 | 2 | 1 | 9 | 3 | 0 | 3 | 5 | 10 | 1 | 4 | 3 | X |  | XXX | XXXX |
| 19 | 58 | 145 | 117 | 2 | 0 | 0 | 38 | 14 | 0 | 2 | 2 | 8 | 3 | 14 | 10 |  |  | XXX | XXXX |
| 2 | 1 | 4 | 0 | 2 | 0 | 1 | 0 | 0 | 0 | 48 | 164 | 126 | 0 | 1 | 0 |  |  | XXX |  |
| 20 | 1 | 8 | 18 | 11 | 2 | 5 | 4 | 4 | 1 | 0 | 0 | 2 | 0 | 1 | 3 | X |  | XXX |  |
| 21 | 8 | 67 | 55 | 3 | 0 | 0 | 8 | 3 | 0 | 0 | 3 | 15 | 0 | 3 | 6 |  |  | XXX | XXXX |
| 22 | 7 | 14 | 6 | 3 | 2 | 2 | 0 | 0 | 0 | 34 | 36 | 8 | 1 | 2 | 2 | X |  |  |  |
| 23/DV6 | 1 | 5 | 0 | 2 | 0 | 0 | 246 | 123 | 23 | 0 | 1 | 1 | 0 | 0 | 1 |  |  |  |  |
| 24 | 2 | 1 | 0 | 6 | 0 | 1 | 27 | 5 | 1 | 3 | 2 | 3 | 2 | 6 | 4 |  |  |  |  |
| 25 | 1 | 3 | 2 | 44 | 55 | 40 | 12 | 6 | 0 | 7 | 17 | 6 | 0 | 0 | 0 |  |  |  |  |
| 26-1 | 4 | 6 | 3 | 3 | 0 | 1 | 112 | 55 | 1 | 2 | 1 | 3 | 0 | 0 | 0 |  |  |  |  |
| 26-2 | 0 | 1 | 1 | 2 | 0 | 0 | 7 | 8 | 2 | 4 | 7 | 0 | 5 | 20 | 8 |  |  | XXX |  |
| 27 | 0 | 1 | 0 | 13 | 0 | 0 | 5 | 1 | 0 | 2 | 14 | 2 | 1 | 0 | 0 |  |  |  |  |
| 29/DV5 | 4 | 8 | 10 | 67 | 92 | 36 | 14 | 0 | 0 | 4 | 8 | 3 | 3 | 2 | 5 | X |  | XXX |  |
| 3 | 3 | 11 | 2 | 1 | 0 | 0 | 24 | 11 | 33 | 1 | 11 | 5 | 24 | 157 | 77 | X | XX | XXX | XXXX |
| 30 | 0 | 1 | 1 | 1 | 0 | 0 | 0 | 1 | 1 | 0 | 0 | 0 | 0 | 0 | 0 |  |  |  |  |
| 34 | 0 | 0 | 0 | 1 | 0 | 0 | 0 | 0 | 0 | 0 | 0 | 0 | 0 | 0 | 0 |  |  |  |  |
| 35 | 0 | 0 | 0 | 3 | 0 | 0 | 3 | 0 | 2 | 0 | 1 | 1 | 0 | 8 | 4 |  |  |  |  |
| 36/DV7 | 0 | 2 | 0 | 0 | 0 | 0 | 3 | 1 | 0 | 2 | 2 | 2 | 1 | 1 | 3 |  |  |  |  |
| 38-1 | 0 | 1 | 0 | 4 | 0 | 0 | 5 | 4 | 0 | 0 | 1 | 0 | 0 | 0 | 0 |  |  |  |  |
| 38-2/DV8 | 0 | 0 | 0 | 38 | 51 | 21 | 9 | 10 | 0 | 0 | 6 | 2 | 0 | 22 | 16 |  |  | XXX |  |
| 39 | 10 | 9 | 1 | 0 | 0 | 1 | 93 | 46 | 21 | 0 | 0 | 2 | 9 | 43 | 37 |  |  | XXX |  |
| 4 | 1 | 0 | 0 | 1 | 0 | 0 | 2 | 0 | 1 | 0 | 1 | 3 | 0 | 1 | 1 |  |  |  |  |
| 40 | 0 | 0 | 1 | 0 | 0 | 0 | 0 | 0 | 0 | 0 | 0 | 0 | 0 | 0 | 1 |  |  |  |  |
| 41 | 1 | 4 | 1 | 6 | 0 | 0 | 5 | 2 | 0 | 0 | 4 | 1 | 0 | 5 | 4 |  |  |  |  |
| 5 | 4 | 7 | 3 | 5 | 0 | 1 | 182 | 60 | 15 | 3 | 9 | 6 | 1 | 4 | 1 | X |  | XXX |  |
| 6 | 7 | 21 | 14 | 0 | 0 | 0 | 7 | 0 | 1 | 9 | 12 | 0 | 0 | 0 | 1 |  |  | XXX |  |
| 8-1 | 5 | 12 | 13 | 0 | 0 | 0 | 2 | 3 | 1 | 0 | 1 | 0 | 0 | 1 | 3 |  |  | XXX |  |
| 8-2 | 1 | 9 | 0 | 2 | 1 | 1 | 3 | 2 | 0 | 0 | 1 | 3 | 0 | 2 | 0 |  |  |  |  |
| 8-3 | 1 | 10 | 2 | 1 | 0 | 1 | 4 | 1 | 1 | 107 | 97 | 34 | 16 | 40 | 29 | X |  | XXX |  |
| 8-4 | 2 | 0 | 0 | 0 | 0 | 0 | 6 | 2 | 0 | 2 | 3 | 0 | 1 | 2 | 1 |  |  |  |  |
| 8-6 | 2 | 3 | 2 | 15 | 40 | 22 | 0 | 1 | 2 | 1 | 2 | 7 | 0 | 1 | 0 | X |  |  |  |
| 9-2 | 11 | 6 | 2 | 3 | 2 | 2 | 11 | 3 | 1 | 0 | 0 | 4 | 0 | 0 | 0 |  |  |  |  |

X = at least 4 donors with reactivities to both peptides
XX = at least 3 donors with 5 clones reacting to both peptides
XXX = at least 1 donor with 5 clones reacting to both peptides and increased number of reacting clones vs. UNT
XXXX = 2 or more donors with 5 clones reacting to both peptides and increased number of reacting clones vs. UNT

**TABLE 2.**
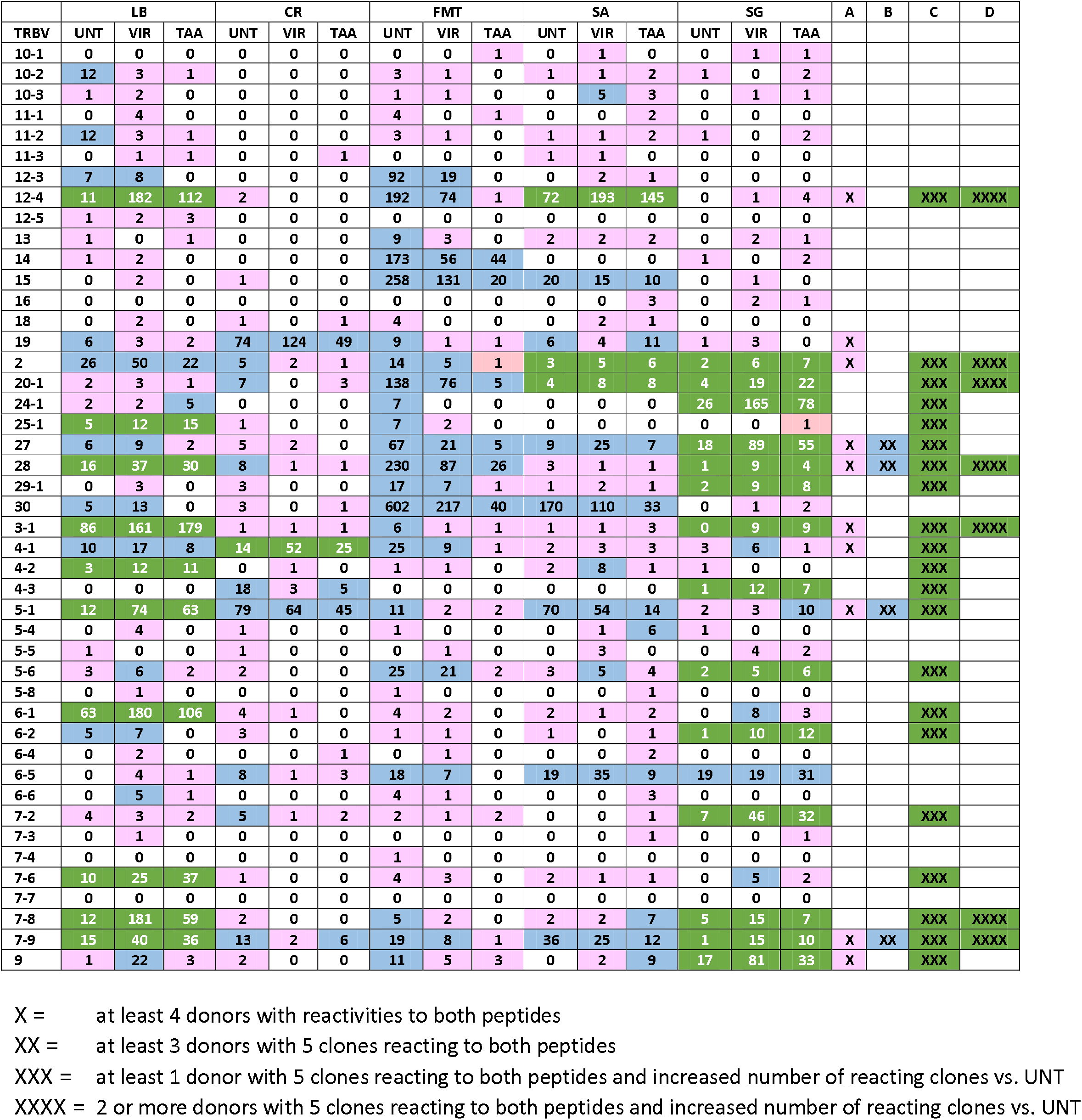
Number of TRBV clonotypes identified in each donor.

### TRAV & TRBV interactions elicited by VIR and TAA antigens

The subsequent analysis was focused on the single cells in the CD8^+^ T_TE_ subset expanded in culture *ex vivo* by treatment with the VIR or TAA antigens. Such results were obtained assessing the co-expression of the TRAV and TRBV molecules in the same cell clone. Overall, selected TRAV/TRBV combinations were significantly expanded after induction with both VIR and TAA antigens, compared to the untreated samples. In particular, these are TRAV3/TRBV24.1, TRAV19/TRBV6.1, TRAV2/TRBV12.4, TRAV17/TRBV12.4, TRAV12.3/TRBV3.1 (Fig. 4). Only the combination TRAV19/TRBV6.1 has been previously reported as TCR SB27 and CA5, binding the HLA-B^*^3508-restricted EBV LPEPLPQGQLTAY peptide^21-23^. All others are not found in publicly searchable databases.

**Fig. 4.**
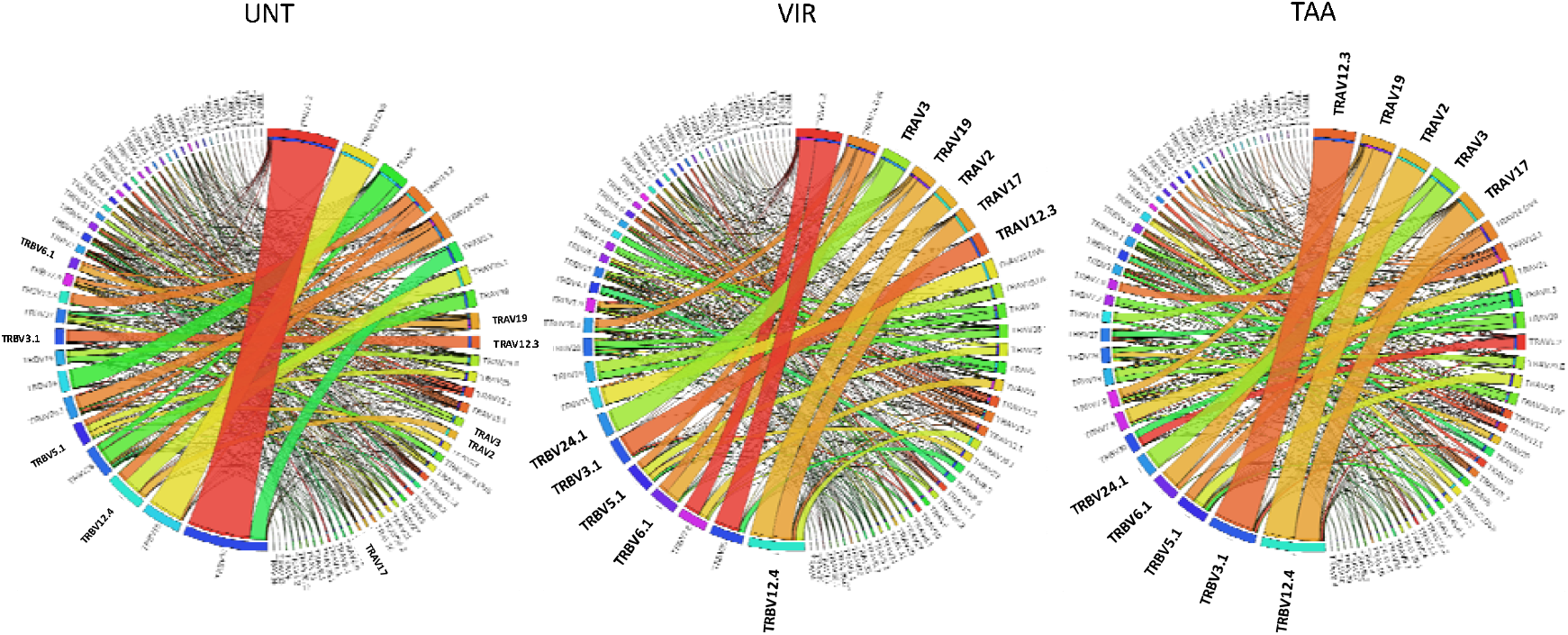
TRAV and TRBV pairing. Circos plots of TRAV and TRBV clonotype pairing in samples from all the donors with the three treatments. Outer arch segment colored by TRAV and TRBV usage. TRAV–TRBV gene pairing indicated by connecting lines, which are colored based on their TRAV usage and segmented, based on their CRD3α and CDR3β sequence. The thickness is proportional to TCR clone number with the respective pair. UNT, untreated; VIR, viral antigen; TAA, tumor antigen.

The CD8^+^ T_TE_ clones expanded by both VIR and TAA antigens show 2 distinct patterns of TRAV/TRBV combinations. As an example, **pattern 1** is characterized by a single TRAV/TRBV interaction in all clones expanded after both treatment (e.g. TRAV3/TRBV24-1); **pattern 2** is characterized by multiple interactions between 1 TRAV and different TRBV molecules in the clones expanded after both treatment (Fig. 5 A-B). Interestingly, the same patterns are followed by the clones identified in the corresponding untreated samples.

**Fig. 5.**
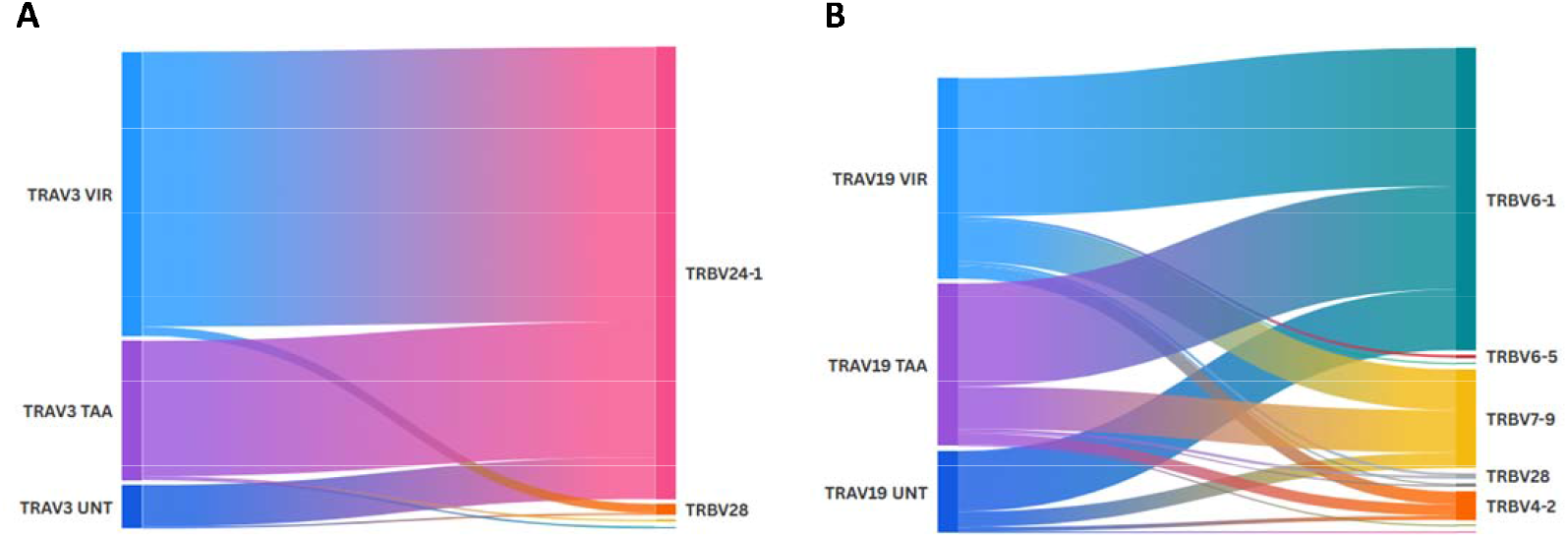
TRAV and TRBV paring pattern. Different patterns of TRAV and TRBV clonotype pairing in samples after the three treatments. Examples of single (A) and multiple (B) pairing are shown in the Sankey diagrams.

### Full TCR combinations elicited by VIR and TAA antigens

In order to evaluate the full CDR3αβ regions involved in the direct interaction between the TCR and the epitope, the TRAV/TRAJ - TRBV/TRBJ combinations were assessed for each CD8^+^ T_TE_ clone expanded by both VIR and TAA antigens. Emphasis was placed on the J (Joining) regions, which provide the primary structural diversity for antigen recognition. The results showed that the prevalent TRAJ and TRBJ regions in the CD8^+^ T_TE_ clones expanded by both VIR and TAA antigens are the same. The only exceptions are the TRBJ1-1 and the TRBJ2-2, which are prevalent only in the clones expanded by the VIR or the TAA antigen, respectively (Fig. 6A and B).

**Fig. 6.**
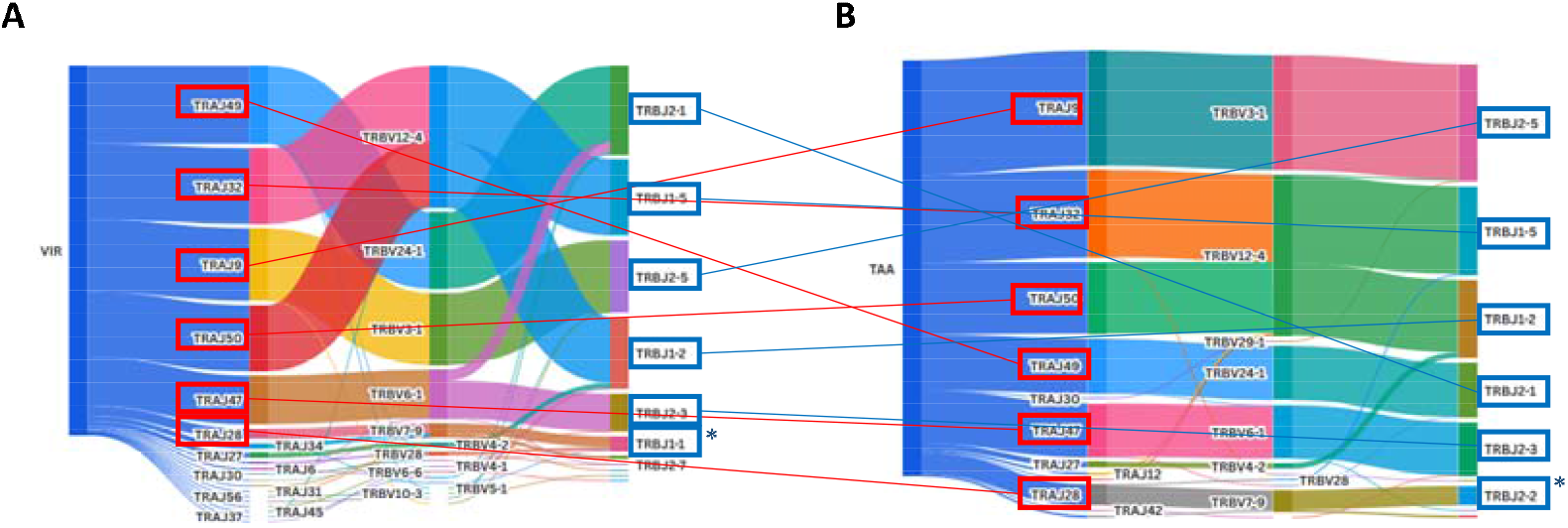
Junction (J) regions in TCRs expanded by VIR and TAA antigens. Junction (J) regions pairing with variable (V) regions are common to TCRs expanded in all the donors by both VIR (A) and TAA (B) epitopes.

Looking into each CD8^+^ T_TE_ clone expanded by both VIR and TAA antigens, the results show that each CDR3αβ region is made of a single (or essentially single) TRAV/TRAJ/TRBV/TRBJ combination. The only exception are the clones expressing the TRAV19, which is found combined with multiple TRAJ/TRBV/TRBJ molecules (Fig. 7)

**Fig. 7.**
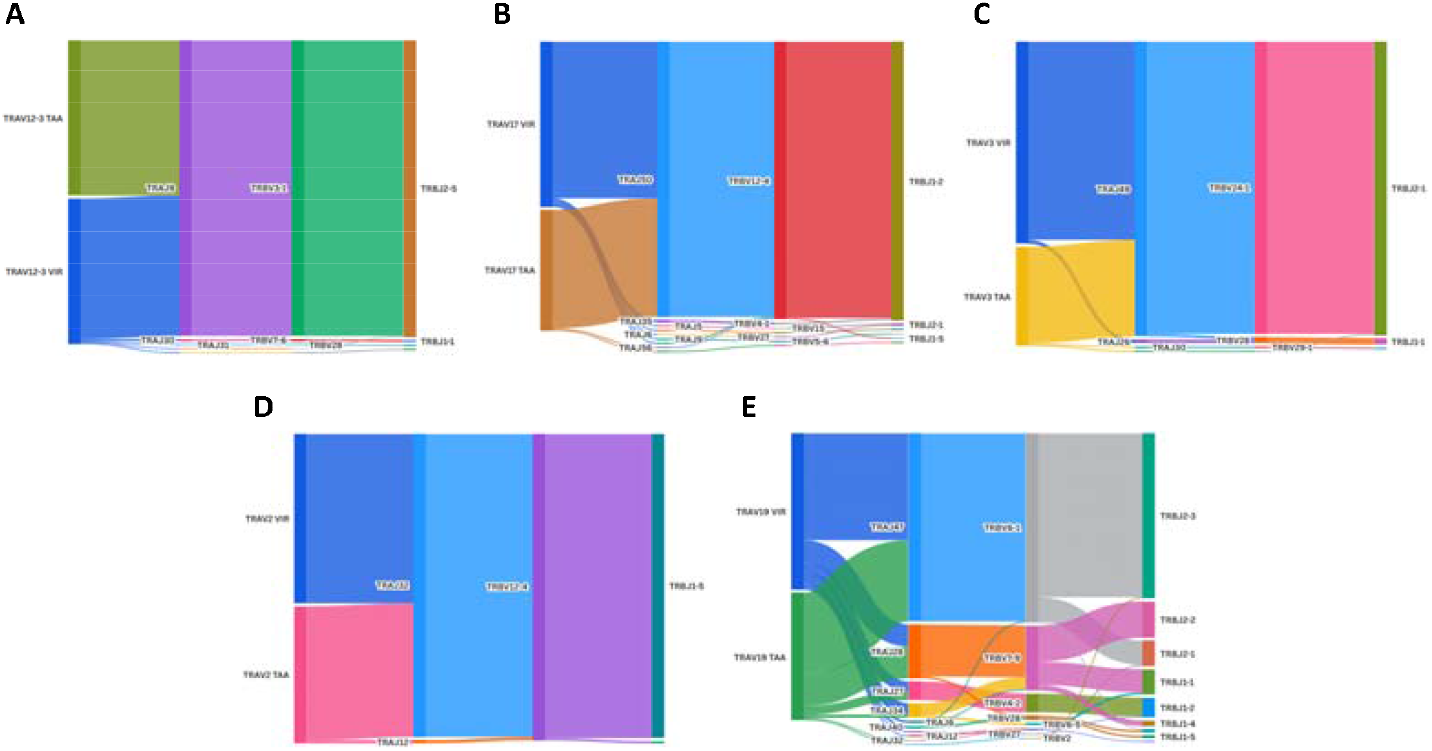
Full TCR pairing expanded by VIR and TAA antigens. Different patterns of TCRs clonotype pairing in samples expanded by both VIR and TAA epitopes. Single (A-D) and multiple (E) pairing are shown in the Sankey diagrams.

### CDR3αβ motifs associated to induction by VIR and TAA antigens

The CDR3αβ motifs of CD8^+^ T_TE_ clones expanded after induction with both VIR and TAA were derived. They are an expansion of motifs found also in the UNT sample with few exceptions (i.e. Trav17 Traj50 Trbv12-4 Trbj1-2), which is a *de novo* identification only in clones expanded by the VIR and TAA antigens (TABLE 3). In support of the biological relevance of such findings, a number of the same clones are identified in the CD8^+^ actively replicating subset, supporting the evidence that they have been activated by the treatment with both peptides (TABLE 3).

**TABLE 3.**
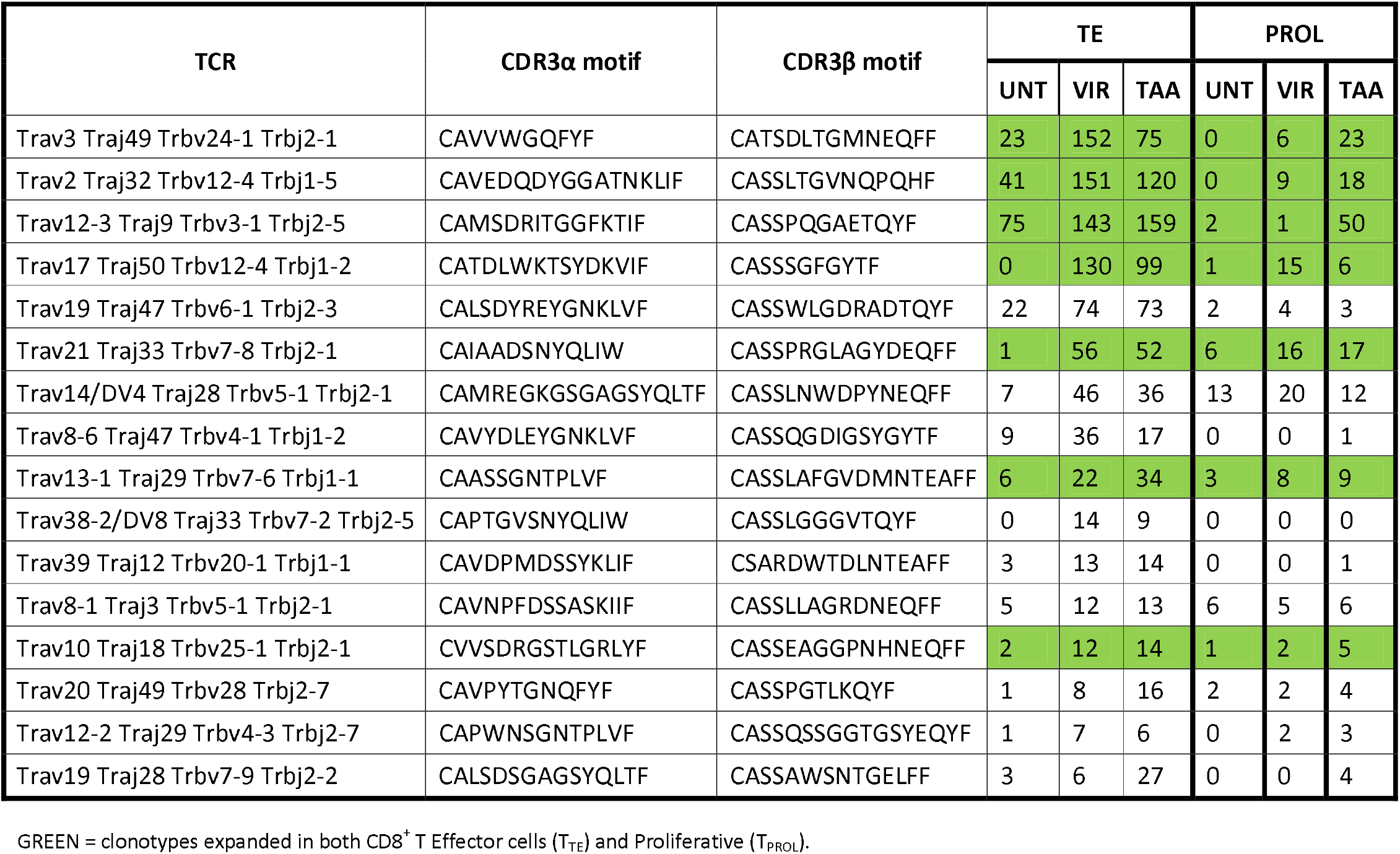
CDR3αβ motifs identified in the TRAV/TRBV clonotypes expanded in CD8^+^ T cells by treatment with VIR and TAA.

The publicly available databases were interrogated to verify whether such combinations have been already reported by others. The analysis showed that the T cell clones characterized by TRAV/TRAJ or the TRBV/TRBJ combinations found in the present study, have been independently previously reported in literature. On the contrary, clones with exactly the same full TRAV/TRAJ/TRBV/TRBJ combination have never been reported before. Moreover, the exact CDR3α and CDR3β motifs expanded with both VIR and TAA antigens have never been reported before. The latter were piled up with sequences from the literature corresponding to the same TRAV/TRAJ or the TRBV/TRBJ combinations and seq logos were generated. For each motif, highly conserved residues are found at both the N-terminus (V region) and the C-terminus (J region), with different degrees of variability is observed in the central regions (Fig. 8).

**Fig. 8.**
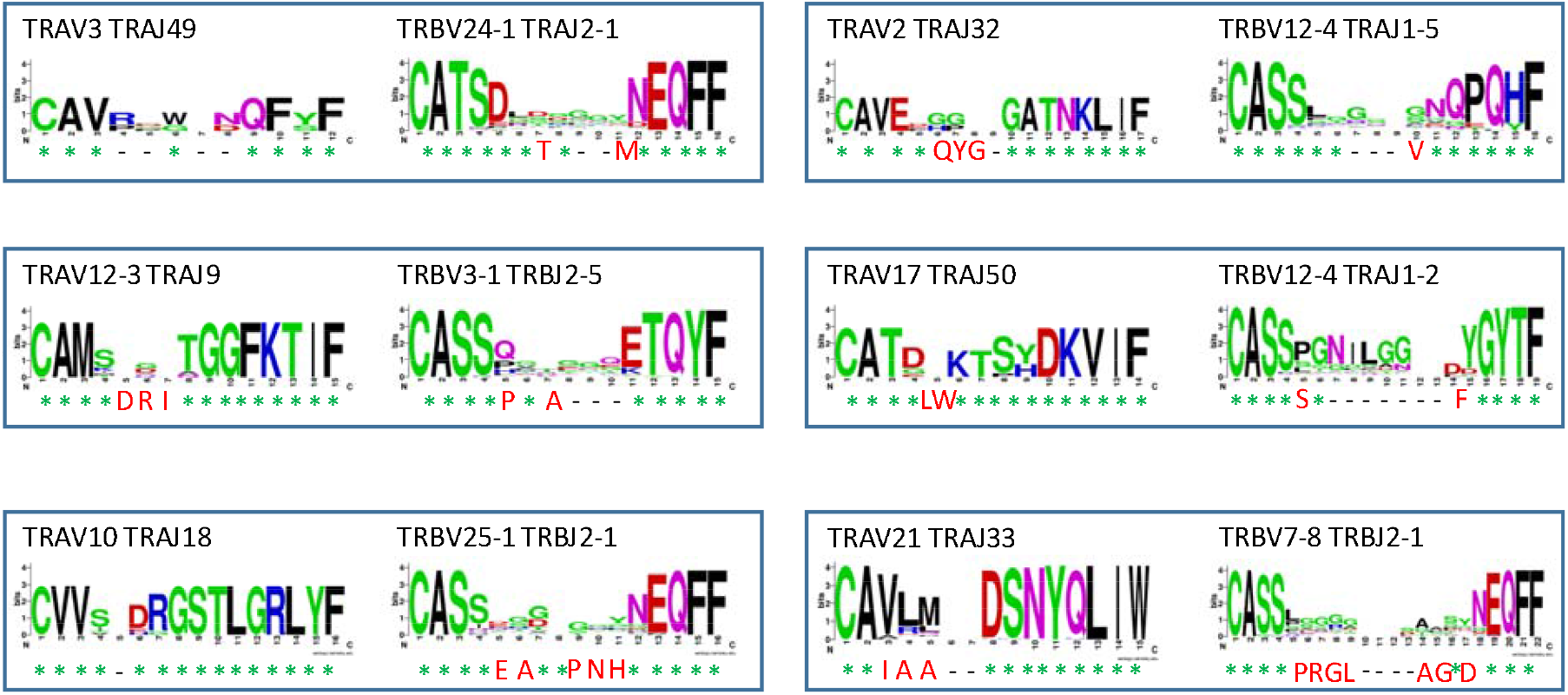
CDR3αβ motifs TCR expanded by VIR and TAA antigens. Consensus logo representations of matched paired CDR3α and CDR3β sequence motifs from publicly available database, corresponding to those identified in the study. The height of the letters indicates the frequency of the specific amino acid residue in that specific position of the epitope. At the bottom the motifs identified in the study are reported. Asterisk = identical residue; red color letter = different residue; hyphen = delition.

### Structural modeling of TCR-pMHC class I complexes

The three-dimensional structure of complexes including the TCR CDR1, CDR2 and CDR3 αβ sequences amplified in the samples after treatment with the VIR or the TAA antigens were predicted using comparative modelling (Table 4). Only for the peptides bound to the MM2 TCR the prediction was not returned.

**TABLE 4.**
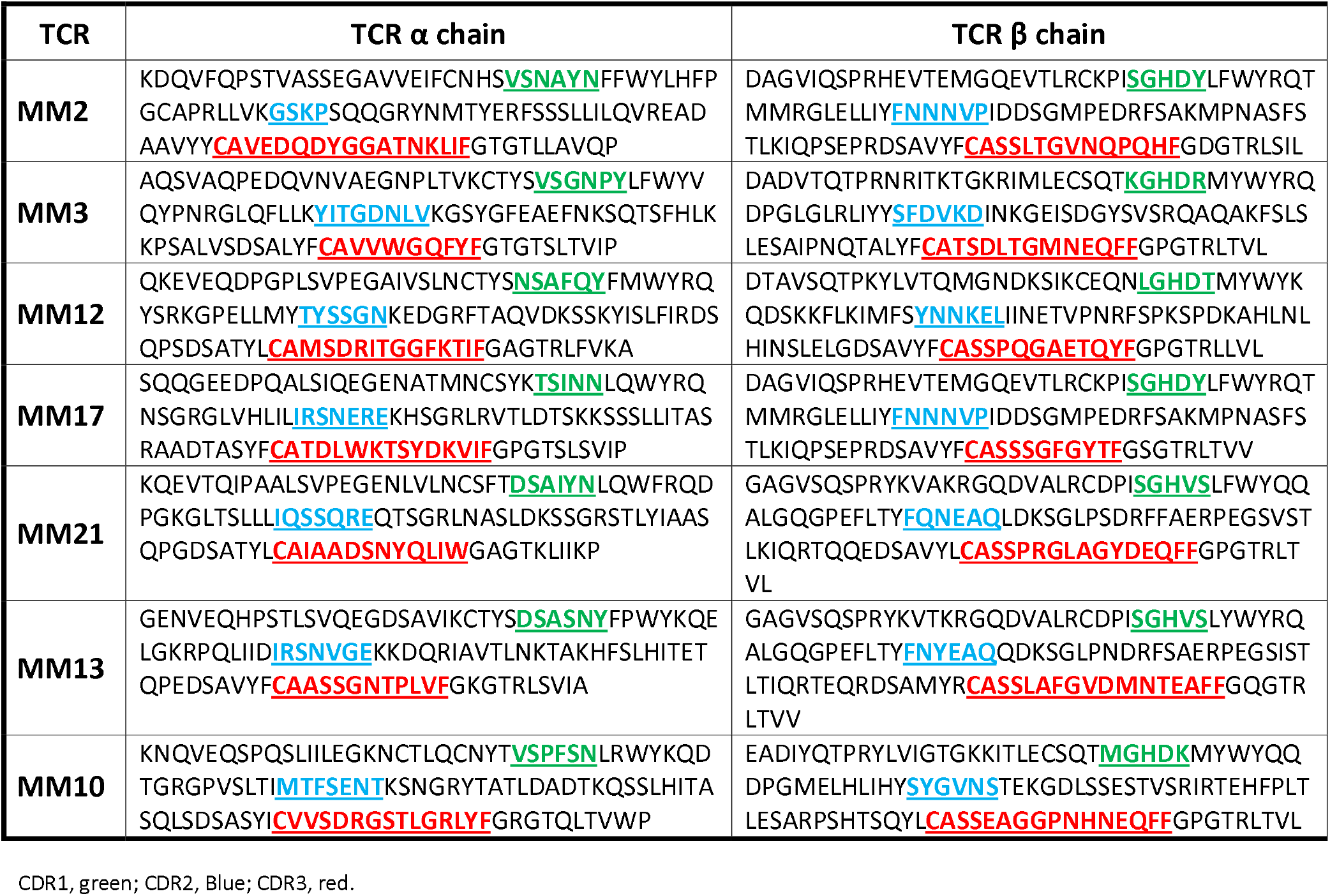
Full TCR αβ chains expanded in CD8^+^ T cells by treatment with VIR and TAA in both CD8^+^ T Effector cells (T_TE_) and Proliferative (T_PROL_)

The predicted conformations of the VIR and the TAA peptides, derived from the binding with all the TCR αβ chains sequences identified in the present study, may be perfectly or partially overlapping. Indeed, the residues facing the TCR (p1, p4, p5 and p8) show a high degree of overlapping induced by the interaction with the αβ chains. The residues facing the HLA groove (p2 and p9) show a perfect overlapping (Fig. 9A).

**Fig. 9A.**
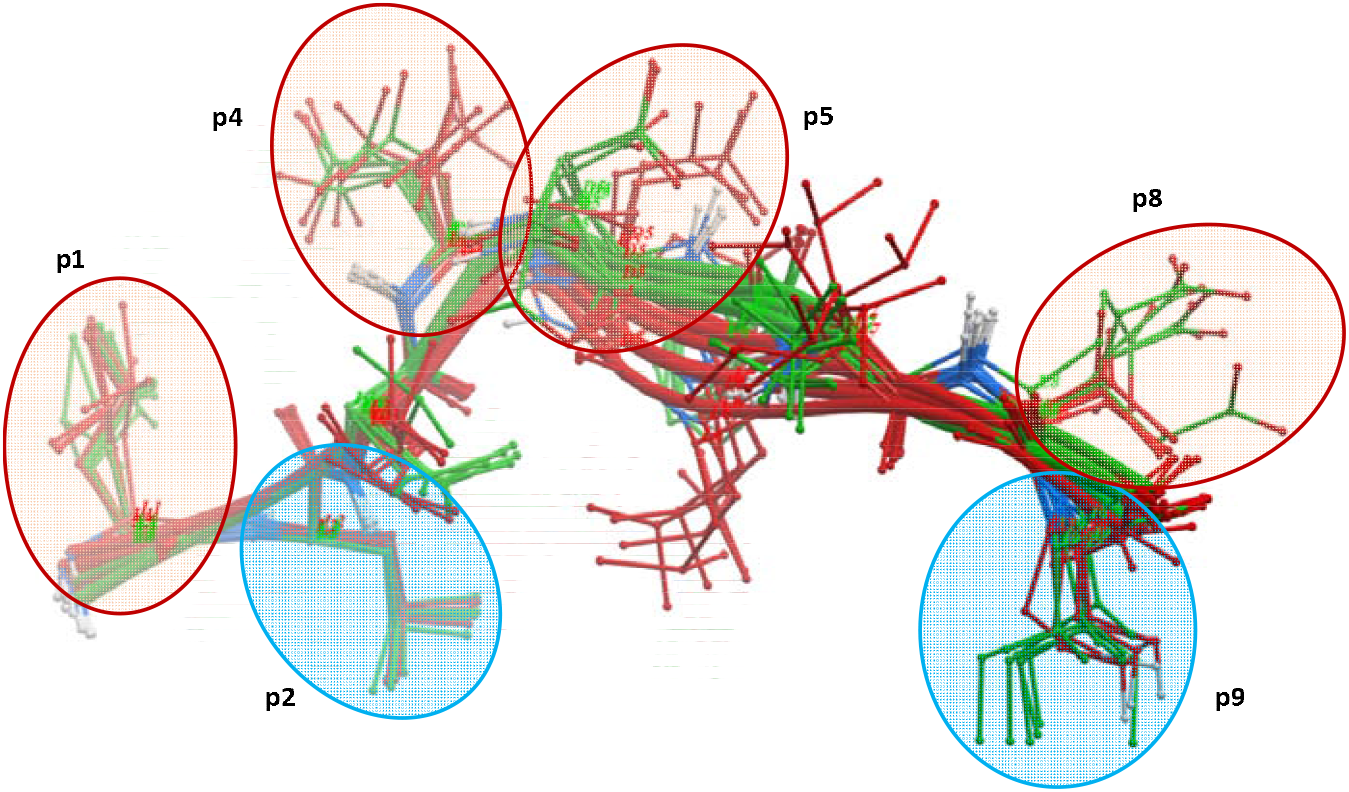
Predicted conformation of VIR and TAA antigens bound to TCRs. The SARS-CoV-2 LLLDDFVEI (green) and the PRDX5 TAA LLLDDLLVS (red) peptides are piled up in their predicted conformation when bound to the different T cell clones described in the text.

The pattern of peptide conformations is echoed in the predicted conformations of the TCRs when bound to the VIR or TAA peptide. Indeed, the MM10, MM12 and MM21 TCRs show a high degree of overlapping, while the MM3, MM13 and MM17 show significant differences (Fig. 9B).

**Fig. 9B.**
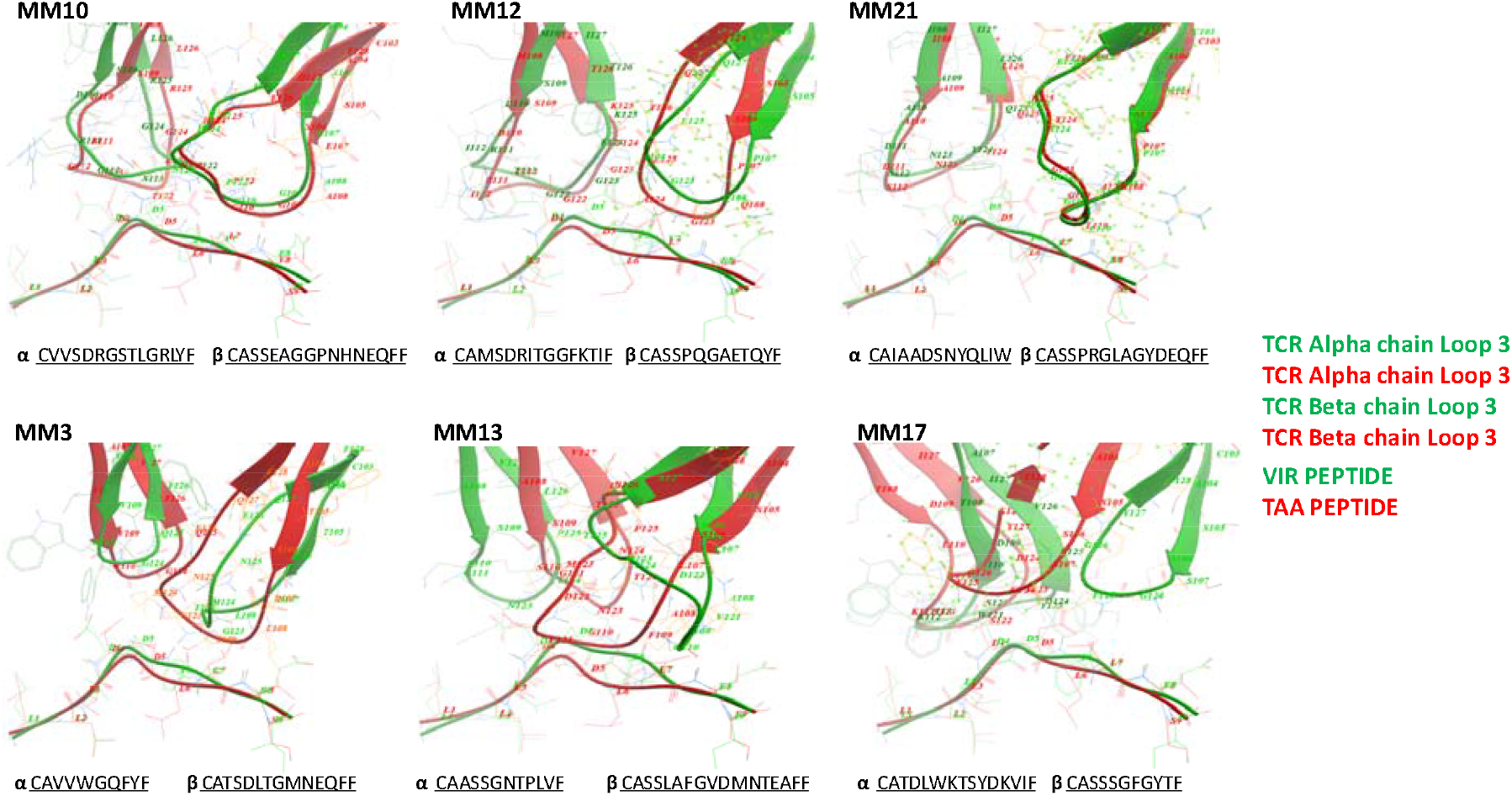
Predicted conformation of TCRs bound to VIR and TAA antigens. The individual CDR3 αβ regions of TCRs are predicted when bound to the VIR (green) or the TAA (red) peptides. Linear amino acid sequences are indicated at the bottom of each image.

## DISCUSSION

In the present study, we analyzed the TCR αβ sequences of CD8^+^ T cell clones induced by the paired SARS-CoV-2 epitope LLLDDFVEI (VIR) and the PRDX5 tumor associated antigen LLLDDLLVS (TAA). The latter two peptides have previously been described by our group to share sequence and structural similarities (molecular mimicry)^10^. The analysis has been extensively performed on CD8^+^ T Effector cells (T_TE_) which are the subset actively responding to a stimulus^20^.

The repertoire of endogenous T cell receptor (TCR) CDR3 αβ motifs in the subjects included in the analysis shows a private pattern of amplified TRAV and TRBV genes, suggesting the ongoing individual immune CD8^+^ T cell response to external stimuli. On the contrary, the repertoire in the Effector memory (T_EM_) subset shows an about uniform distribution of TRAV and TRBV genes in all subjects, suggesting the broad representation of memory T cell clones elicited along the time.

The individual repertoire in the endogenous pool CD8^+^ T Effector cells (T_TE_) was confirmed in the donor-specific set of TRAV and TRBV clones expanded by the *ex vivo* induction of the VIR or TAA peptide.

The UMAP projection of CD8^+^ T cells from all the 5 donors revealed a consistent pattern of activation induced by the treatments with both the VIR and TAA peptides. The stimulation with both peptides induced a highly dynamic CD8^+^ T-cell landscape characterized by extensive proliferation and phenotypic plasticity. Such a picture was more predominant in the sample treated with the VIR peptide, which could be due to a previous priming of T cells by the SARS-CoV2 antigen. Stimulation with the TAA peptide showed feature more consistent with advanced effector differentiation and reduced proliferative capacity, which need additional functional and transcriptional markers to conclusively confirm a terminal differentiation.

The majority of such clones are expanded by either the VIR or the TAA epitope in a very limited number (<5), which unlikely have a relevant biological meaning. Therefore, considering as relevant only clones expanded by both epitopes in a number >5, 21 TRAV and TRBV are independently identified in at least a single donor and 7 in at least 2 donors. Five TRAV/TRBV combinations co-expressed in the same T_TE_ clones are significantly expanded by *ex vivo* by treatment with the VIR or TAA antigens vs. the untreated samples (UNT). Of these, none has been previously reported, except for the combination TRAV19/TRBV6.1 which has been identified to bind the HLA-B^*^3508-restricted EBV LPEPLPQGQLTAY peptide^21-23^. Furthermore, the full picture of the analysis is provided by including the J regions of CDR3, which is essential for binding the epitope^24^. The results show that, collectively, all the T cell clones expanded by both VIR and TAA peptides are characterized by the same J regions in both α and β chains. However, the different clones show a gradient of complexity going from a single combination (TRAV:TRAJ:TRBV:TRBJ = 1:1:1:1) to a multiple combination (e.g. TRAV:TRAJ:TRBV:TRBJ = 1:4:4:6). This confirms the high degeneracy of T cell response in binding a single epitope, needed for covering a broad spectrum of targets which largely exceed the number of T cells in our human body^25^. Nevertheless, the very same combinations have been identified also in the profilerative CD8^+^ T cell subset after treatment with the VIR or the TAA peptides, supporting the notion that both peptides are able to elicit activation and proliferation of cross-reactive CD8^+^ T cells. The exact CDR3αβ aminoacid motifs expanded with both VIR and TAA antigens have never been reported before, however they show high degree of homology with sequences in literature corresponding to the same TRAV/TRAJ/TRBV/TRBJ molecules. This is particularly evident in the J regions of both αβ chains, which are >95% homologous to the published motifs.

Finally, the predicted conformation of the complex made by VIR or TAA peptides when bound to MM10, MM12 and MM21 TCRs show high degree of overlapping.

All the findings in the present study gather evidences showing that a number of cross-reacting TCRs are elicited by similar viral-derived and tumor-associated antigens. This fully support the concept that non-self viral antigens may prime the immune system against several over-expressed self TAAs. In particular, as described in the present study, the HLA-A^*^02:01-restricted SARS-CoV-2 epitope LLLDDFVEI elicits T cells with TCRs cross-reacting with the PRDX5 tumor associated antigen LLLDDLLVS. The implication is that such viral antigen may be used to develop off-the-shelf therapeutic vaccine formulations specific for the melanoma. Furthermore, it strongly suggest that the recent COVID-19 pandemics and/or vaccination may have induced a memory T cell population to prevent melanoma or improving its clinical prognosis.

In conclusion, the present study provides robust proofs to support the development of preventive/therapeutic cancer vaccines based on highly immunogenic non-self microbe-derived antigens for eliciting a strong cross-reacting anti-cancer T cell response. This could likely represent the definitive solution for the poor efficacy of the cancer vaccines based on over-expressed self TAAs and boosting a second-wave of off-the-shelf vaccines.

## Acknowledgements

Not applicable.

## Declarations

### 1. Ethics approval and consent to participate

The study has been approved by the Institutional Ethical Committee of the Istituto Nazionale Tumori - IRCCS - “Fond G. Pascale” – NAPOLI IT (Protocol n. 57/22 OSS “HepAnt – Tumor antigen discovery for innovative cancer immunotherapies in HCC: from benchside to bedside” e subsequent amendments).

### 2. Consent for publication

The corresponding author has received consent for publication.

### 3. Availability of data and material

Data and material will be deposited and publicly available.

### 4. Competing interests

The authors declare no potential conflicts of interest.

### 5. Funding

The study was funded by the Italian Ministry of Health through Institutional “Ricerca Corrente” (Projects L2/30_25 to LB; L2/35_25 to MT; L4/48_25 to CR); the PNRR Ministero Salute PNRR-POC-2022-12375769 “Molecular mimicry to improve liver cancer immunotherapy” (2023 – 2025) (to LB).

### 6. Authors’ contributions

CR performed 40% of all the antigen prediction analyses; BC and AM performed the remaining 20% (10% each) of the antigen prediction analyses. SM and BC performed the PBMCs ex vivo stimulations and RNA extractions. NC performed all the bibliographic searches. MT and LB designed the structure of the review article, supervised the analysis and drafted the manuscript.

## Notes

### Competing Interest Statement

The authors have declared no competing interest.

## REFERENCES

1. Lin MJ, Svensson-Arvelund J, Lubitz GS, Marabelle A, Melero I, Brown BD, Brody JD. Cancer vaccines: the next immunotherapy frontier. Nat Cancer. 2022 Aug;3(8):911–926. doi: 10.1038/s43018-022-00418-6.

2. Katsikis PD, Ishii KJ, Schliehe C. Challenges in developing personalized neoantigen cancer vaccines. Nat Rev Immunol. 2024 Mar;24(3):213–227. doi: 10.1038/s41577-023-00937-y. Epub 2023 Oct 2.

3. Buonaguro L, Cerullo V. Pathogens: Our Allies against Cancer? Mol Ther. 2021 Jan 6;29(1):10–12. doi: 10.1016/j.ymthe.2020.12.005.

4. Tagliamonte M, Cavalluzzo B, Mauriello A, Ragone C, Buonaguro FM, Tornesello ML, Buonaguro L. Molecular mimicry and cancer vaccine development. Mol Cancer. 2023 Apr 26;22(1):75. doi: 10.1186/s12943-023-01776-0.

5. Sewell, A. Why must T cells be cross-reactive?. Nat Rev Immunol 12, 669–677 (2012). 10.1038/nri3279;

6. Ragone C, Manolio C, Cavalluzzo B, Mauriello A, Tornesello ML, Buonaguro FM, Castiglione F, Vitagliano L, Iaccarino E, Ruvo M, Tagliamonte M, Buonaguro L. Identification and validation of viral antigens sharing sequence and structural homology with tumor-associated antigens (TAAs). J Immunother Cancer. 2021 May;9(5):e002694. doi: 10.1136/jitc-2021-002694.

7. Pearson JRD, Puig-Saenz C, Thomas JE, Hardowar LD, Ahmad M, Wainwright LC, McVicar AM, Brentville VA, Tinsley CJ, Pockley AG, Durrant LG, McArdle SEB. TRP-2 / gp100 DNA vaccine and PD-1 checkpoint blockade combination for the treatment of intracranial tumors. Cancer Immunol Immunother. 2024 Jul 2;73(9):178. doi: 10.1007/s00262-024-03770-x.

8. Suda T, Tsunoda T, Uchida N, Watanabe T, Hasegawa S, Satoh S, Ohgi S, Furukawa Y, Nakamura Y, Tahara H. Identification of secernin 1 as a novel immunotherapy target for gastric cancer using the expression profiles of cDNA microarray. Cancer Sci. 2006 May;97(5):411–9. doi: 10.1111/j.1349-7006.2006.00194.x.

9. Ragone C, Manolio C, Mauriello A, Cavalluzzo B, Buonaguro FM, Tornesello ML, Tagliamonte M, Buonaguro L. Molecular mimicry between tumor associated antigens and microbiota-derived epitopes. J Transl Med. 2022 Jul 14;20(1):316. doi: 10.1186/s12967-022-03512-6.

10. Ragone C, Mauriello A, Cavalluzzo B, Cavalcanti E, Russo L, Manolio C, Mangano S, Cembrola B, Tagliamonte M, Buonaguro L. Molecular mimicry of SARS-COV-2 antigens as a possible natural anti-cancer preventive immunization. Front Immunol. 2024 Jun 14;15:1398002. doi: 10.3389/fimmu.2024.1398002.

11. Vujanovic L, Shi J, Kirkwood JM, Storkus WJ, Butterfield LH. Molecular mimicry of MAGE-A6 and Mycoplasma penetrans HF-2 epitopes in the induction of antitumor CD8+ T-cell responses. Oncoimmunology. 2014 Nov 14;3(8):e954501. doi: 10.4161/21624011.2014.954501.

12. Petrizzo A, Tagliamonte M, Mauriello A, Costa V, Aprile M, Esposito R, Caporale A, Luciano A, Arra C, Tornesello ML, Buonaguro FM, Buonaguro L. Unique true predicted neoantigens (TPNAs) correlates with anti-tumor immune control in HCC patients. J Transl Med. 2018 Oct 19;16(1):286. doi: 10.1186/s12967-018-1662-9.

13. Cavalluzzo B, Mauriello A, Ragone C, Manolio C, Tornesello ML, Buonaguro FM, Tvingsholm SA, Hadrup SR, Tagliamonte M, Buonaguro L. Novel Molecular Targets for Hepatocellular Carcinoma. Cancers (Basel). 2021 Dec 28;14(1):140. doi: 10.3390/cancers14010140.

14. Manolio C, Ragone C, Cavalluzzo B, Mauriello A, Tornesello ML, Buonaguro FM, Salomone Megna A, D’Alessio G, Penta R, Tagliamonte M, Buonaguro L. Antigenic molecular mimicry in viral-mediated protection from cancer: the HIV case. J Transl Med. 2022 Oct 15;20(1):472. doi: 10.1186/s12967-022-03681-4.

15. Loftus DJ, Castelli C, Clay TM, Squarcina P, Marincola FM, Nishimura MI, Parmiani G, Appella E, Rivoltini L. Identification of epitope mimics recognized by CTL reactive to the melanoma/melanocyte-derived peptide MART-1(27-35). J Exp Med. 1996 Aug 1;184(2):647–57. doi: 10.1084/jem.184.2.647.

16. Pittet MJ, Valmori D, Dunbar PR, Speiser DE, Liénard D, Lejeune F, Fleischhauer K, Cerundolo V, Cerottini JC, Romero P. High frequencies of naive Melan-A/MART-1-specific CD8(+) T cells in a large proportion of human histocompatibility leukocyte antigen (HLA)-A2 individuals. J Exp Med. 1999 Sep 6;190(5):705–15. doi: 10.1084/jem.190.5.705.

17. Chiou SH, Tseng D, Reuben A, Mallajosyula V, Molina IS, Conley S, Wilhelmy J, McSween AM, Yang X, Nishimiya D, Sinha R, Nabet BY, Wang C, Shrager JB, Berry MF, Backhus L, Lui NS, Wakelee HA, Neal JW, Padda SK, Berry GJ, Delaidelli A, Sorensen PH, Sotillo E, Tran P, Benson JA, Richards R, Labanieh L, Klysz DD, Louis DM, Feldman SA, Diehn M, Weissman IL, Zhang J, Wistuba II, Futreal PA, Heymach JV, Garcia KC, Mackall CL, Davis MM. Global analysis of shared T cell specificities in human non-small cell lung cancer enables HLA inference and antigen discovery. Immunity. 2021 Mar 9;54(3):586-602.e8. doi: 10.1016/j.immuni.2021.02.014.

18. Macandog ADG, Catozzi C, Capone M, Nabinejad A, Nanaware PP, Liu S, Vinjamuri S, Stunnenberg JA, Galiè S, Jodice MG, Montani F, Armanini F, Cassano E, Madonna G, Mallardo D, Mazzi B, Pece S, Tagliamonte M, Vanella V, Barberis M, Ferrucci PF, Blank CU, Bouvier M, Andrews MC, Xu X, Santambrogio L, Segata N, Buonaguro L, Cocorocchio E, Ascierto PA, Manzo T, Nezi L. Longitudinal analysis of the gut microbiota during anti-PD-1 therapy reveals stable microbial features of response in melanoma patients. Cell Host Microbe. 2024 Nov 13;32(11):2004-2018.e9. doi: 10.1016/j.chom.2024.10.006.

19. Ottaiano A, Scala S, D’Alterio C, Trotta A, Bello A, Rea G, Picone C, Santorsola M, Petrillo A, Nasti G. Unexpected tumor reduction in metastatic colorectal cancer patients during SARS-Cov-2 infection. Ther Adv Med Oncol. 2021 Apr 29;13:17588359211011455. doi: 10.1177/17588359211011455.

20. Koh CH, Lee S, Kwak M, Kim BS, Chung Y. CD8 T-cell subsets: heterogeneity, functions, and therapeutic potential. Exp Mol Med. 2023 Nov;55(11):2287–2299. doi: 10.1038/s12276-023-01105-x.

21. Tynan FE, Burrows SR, Buckle AM, Clements CS, Borg NA, Miles JJ, Beddoe T, Whisstock JC, Wilce MC, Silins SL, Burrows JM, Kjer-Nielsen L, Kostenko L, Purcell AW, McCluskey J, Rossjohn J. T cell receptor recognition of a ‘super-bulged’ major histocompatibility complex class I-bound peptide. Nat Immunol. 2005 Nov;6(11):1114–22. doi: 10.1038/ni1257.

22. Burrows SR, Chen Z, Archbold JK, Tynan FE, Beddoe T, Kjer-Nielsen L, Miles JJ, Khanna R, Moss DJ, Liu YC, Gras S, Kostenko L, Brennan RM, Clements CS, Brooks AG, Purcell AW, McCluskey J, Rossjohn J. Hard wiring of T cell receptor specificity for the major histocompatibility complex is underpinned by TCR adaptability. Proc Natl Acad Sci U S A. 2010 Jun 8;107(23):10608–13. doi: 10.1073/pnas.1004926107.

23. Liu YC, Miles JJ, Neller MA, Gostick E, Price DA, Purcell AW, McCluskey J, Burrows SR, Rossjohn J, Gras S. Highly divergent T-cell receptor binding modes underlie specific recognition of a bulged viral peptide bound to a human leukocyte antigen class I molecule. J Biol Chem. 2013 May 31;288(22):15442–54. doi: 10.1074/jbc.M112.447185.

24. Krogsgaard M, Davis MM. How T cells ‘see’ antigen. Nat Immunol. 2005 Mar;6(3):239–45. doi: 10.1038/ni1173.

25. Mason D. A very high level of crossreactivity is an essential feature of the T-cell receptor. Immunol Today. 1998 Sep;19(9):395–404. doi: 10.1016/s0167-5699(98)01299-7.

